# Inducible CRISPRi-based operon silencing and selective *in trans* gene complementation in *Borrelia burgdorferi*

**DOI:** 10.1101/2022.12.05.519228

**Authors:** Bryan T. Murphy, Jacob J. Wiepen, Huan He, Ankita S. Pramanik, Jason M. Peters, Brian Stevenson, Wolfram R. Zückert

## Abstract

To accelerate genetic studies on the Lyme disease pathogen *Borrelia burgdorferi*, we developed an enhanced CRISPR interference (CRISPRi) approach for IPTG-inducible repression of specific *B. burgdorferi* genes. The entire system is encoded on a compact 11-kb shuttle vector plasmid that allows for inducible expression of both the sgRNA module and a non-toxic codon-optimized dCas9 protein. We validated this CRISPRi system by targeting the genes encoding for OspA and OspB, abundant surface lipoproteins co-expressed by a single operon, and FlaB, the major subunit forming the periplasmic flagella. As in other systems, sgRNAs complementary to the non-template strand were consistently effective in gene repression, with 4- to 994-fold reductions in targeted transcript levels and concomitant reductions of in proteins levels. Furthermore, we showed that *ospAB* knockdowns could be selectively complemented *in trans* for OspA expression via the insertion of synonymous or non-synonymous CRISPRi-resistant PAM mutant (PAM*) *ospA* alleles into a unique site within the plasmid. Together, this establishes CRISPRi PAM* as a robust new genetic tool to simplify the study of *B. burgdorferi* genes, bypassing the need for gene disruptions by allelic exchange and avoiding rare-codon toxicity from heterologous expression of dCas9.

**SIGNIFICANCE:** *Borrelia burgdorferi*, the causative agent of Lyme disease, is a tick-borne pathogen of global importance. Here, we expand the genetic toolbox for studying *B. burgdorferi* physiology and pathogenesis by establishing a single-plasmid-based CRISPRi system with optional *in trans* complementation for the functional study of essential and non-essential proteins.

## INTRODUCTION

CRISPR-Cas systems naturally function for prokaryotic adaptive immunity against viruses and plasmids, but their synthetic applications continue to revolutionize medicine and research (1). The chromosomal loci encoding these systems contain arrays of short direct repeats separated by spacer sequences of foreign origin. This array is transcribed and processed into CRISPR RNAs which associate with RNA-guided Cas nucleases to locate and cleave complementary nucleic acid sequences (*i.e*., protospacers) in the invading genome of a virus or plasmid. To prevent the spacer from recognizing and cleaving its complement in the chromosome array, Cas nucleases must also recognize a protospacer-adjacent motif (PAM) immediately adjacent to the protospacer sequence on the non-targeted DNA strand. Class 2 CRISPR-Cas systems are particularly unique for their single multidomain effectors (Cas9, Cas12, and Cas13) which are functionally analogous to entire complexes of Cas proteins in Class 1 systems (2). In 2013, Qi and colleagues co-expressed a catalytically dead version of Cas9 (dCas9) from *Streptococcus pyogenes* with customizable single guide RNAs (sgRNAs) that sterically block transcription elongation of the 20-nucleotide protospacer targets with short 5’-NGG-3’ PAM sites (3). Owing to the technique’s superiority for simple, titratable, target-specific, and reversible gene silencing, this CRISPR interference (CRISPRi) method has since become a widely used technique for gene function studies across all three domains of life (3–5), including diverse bacterial species (6–10).

*Borrelia burgdorferi*, the causative spirochete of Lyme disease (11), is evolutionarily distant from other tractable bacterial models (12). It is among the most genetically complex bacteria known with a distinctly segmented genome consisting of a megabase chromosome and large set of circular and linear plasmids (13, 14). Nonetheless, a robust genetic toolbox has been developed to investigate *B. burgdorferi* physiology and pathogenesis (15, 16). This includes (i) a growing number of antibiotic resistance cassettes (17–21) that can be used for gene disruption and complementation by homologous recombination, (ii) *E. coli/B. burgdorferi* shuttle vector plasmids (17, 22, 23), and (iii) hybrid *lac* or *tet* promoters for conditional gene expression (24–26). With these tools, functional assessment of essential genes classically requires the generation of merodiploid *lac*- or *tet*-promoter-driven overexpression intermediates that are then further mutated to disrupt the endogenous gene copy. Here, we developed a fully inducible, non-toxic CRISPRi system with optional *in trans* gene complementation for use in *Borrelia burgdorferi*. We validated this system by targeting a polycistronic operon encoding OspA and OspB, two abundant and well-studied borrelial outer surface lipoproteins that play important roles in the pathogen’s enzootic cycle (27–32).

## RESULTS

Like most bacterial phyla, spirochaetes are predicted to contain native CRISPR-Cas systems based on extensive computational analyses (2). However, Cas homologs found in various *Treponema* and *Leptospira* species are not detectable in the *Borrelia* genus with domain-based NCBI BLAST searches (33). This suggested that synthetic CRISPR systems can be successfully implemented in *B. burgdorferi* and other related species without any prior genomic alterations. To construct a CRISPRi system for *B. burgdorferi*, we modified the Mobile-CRISPRi system described by Peters *et al*. for diverse bacteria (34). We aimed (i) to simplify the experimental workflow of generating knockdown mutants by providing all necessary elements on a single customizable plasmid, (ii) to eliminate the potential for rare-codon usage toxicity in a low GC content organism by codon-optimizing any heterologous proteins, and (iii) to increase stringency of the system by placing both dCas9 and sgRNA CRISPRi modules under inducible promoters. Briefly, in two stepwise NEBuilder HiFi assemblies, *Streptococcus pyogenes-derived* dCas9-myc (SpydCas9-myc) and *trc*-driven sgRNA modules from the Mobile-CRISPRi system were amplified by PCR and combined with a third module containing the backbone of pJSB142, a shuttle vector for isopropyl-β-D-thiogalactopyranoside (IPTG)-inducible expression of genes under a hybrid T5/*lac* promoter (*P_QE30_*) in *B. burgdorferi* (25). *B. burgdorferi* transformants carrying the resulting recombinant plasmid pJJW100 showed a severe growth defect under inducing conditions (not shown), which suggested rare-codon toxicity (35) from SpydCas9. We therefore codon-optimized SpydCas9 using the OPTIMIZER web-based server (Supplementary file 1) (36). The *B. burgdorferi* codon-optimized dCas9 (BbdCas9) nucleotide sequence was synthesized by GenScript and used to replace the original gene in pJJW100 by restriction-ligation, yielding pJJW101 (Fig. 1A). This optimization eliminated any observable toxicity in *B. burgdorferi* (Fig. 1C).

**Figure 1.**
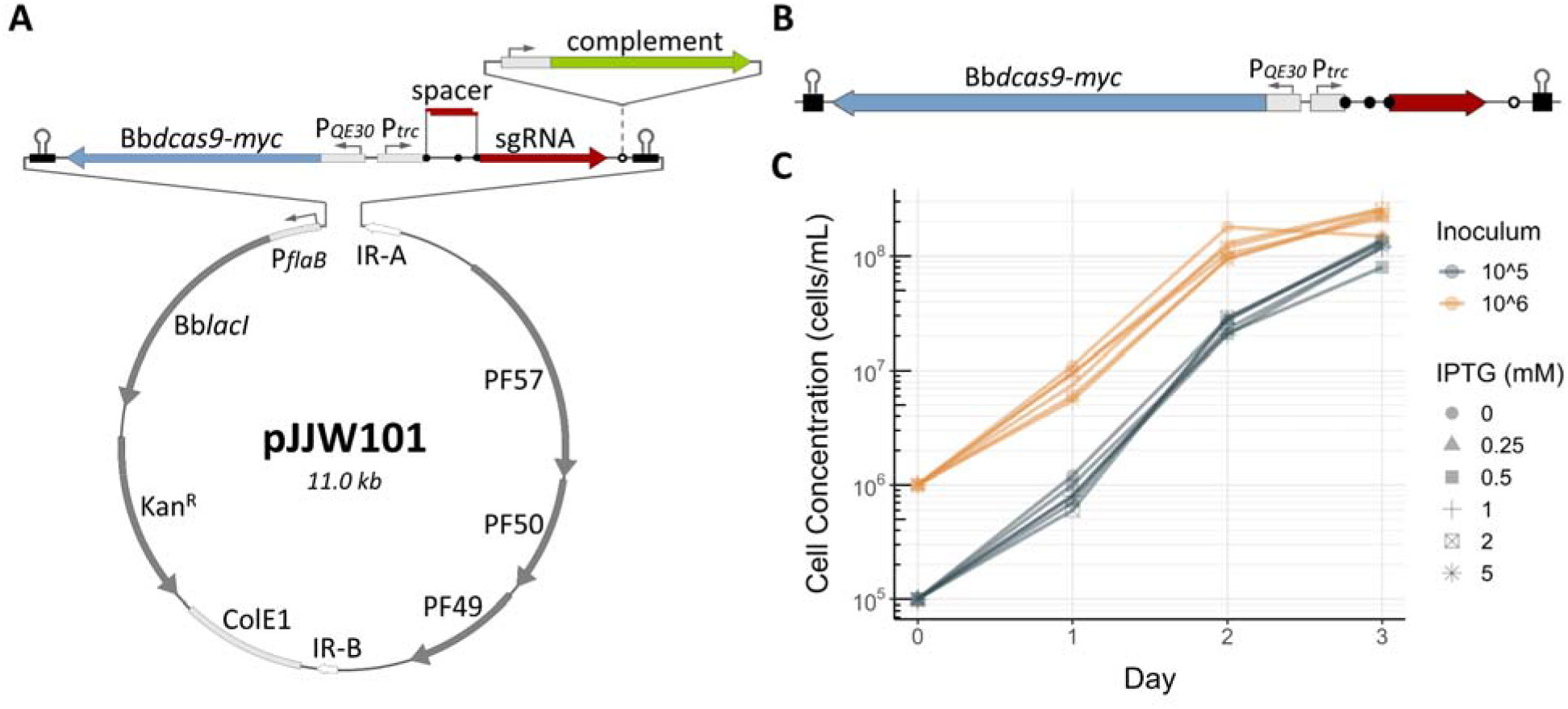
Single-plasmid-based CRISPRi system for *Borrelia burgdorferi*. **(A)** Plasmid map showing the functional elements of pJJW101. pJJW101 contains a pUC (ColE1) high-copy origin of replication and inverted repeat (IR)-flanked *B. burgdorferi* paralogous family (PF) 57, 50 and 49 ORFs which allow for the plasmid’s autonomous replication in *E. coli* and *B. burgdorferi*, respectively (see text). Codon-optimized LacI (BbLacI), constitutively expressed from a *B. burgdorferi flaB* promoter, represses both codon-optimized BbdCas9-myc and sgRNA production under pQE30 (T5/*lac* hybrid) and *trc* (*trp/lac* hybrid) promoters, respectively. Three sequential BsaI sites (black filled circles) are used for the directional insertion of unique gene-targeting spacers in a “scarless” Golden Gate Assembly-like manner that allows digestion and ligation steps to be carried out simultaneously, with the elimination of BsaI recognition sites in the ligated product. A single BglI site (open circle) downstream of the sgRNA module can be used for directional insertion of complementing gene copies or reporters. The plasmid confers resistance to kanamycin (Kan^R^) with the *aph[3’]-IIIa* gene. **(B)** Schematic of the pJJW101 knockdown functional region as an empty vector control without gene-targeting sgRNA spacer insert. **(C)** Liquid culture growth curves of recombinant *B. burgdorferi* B31-e2 carrying pJJW101 (empty vector control). Cultures were inoculated at either 1×10^5^ cells/ml (teal lines) or 1×10^6^ cells/ml (orange lines) and grown in the presence of different IPTG inducer concentrations.

To test the system’s knockdown efficiency, we first designed 20-nucleotide spacers targeting either the non-template (NT) or template (T) strands of *flaB* or *ospA* (Table 2) using the open-source CRISPy-web interface on an artificially concatenated sequence of the *Borrelia burgdorferi* B31 genome (Supplementary file 2) (37). Pairs of complementary primers with 5’-overhangs matching the BsaI cut site sticky ends (Table 1) were synthesized for each chosen spacer sequence and hybridized. The resulting double-stranded oligos were directly ligated into BsaI-cut pJJW101, generating a series of *flaB* and *ospA* knockdown plasmids (Table 4; plasmid IDs 2-10) with simple knockdown functional regions (Fig. 2A). *Borrelia burgdorferi* B31-e2 clones carrying the “empty” pJJW101 vector (lacking any targeting spacer sequence) or one of the nine derived knockdown plasmids were cultured, and their whole cell protein profiles were determined by SDS-PAGE and immunoblotting at 36 hours post-induction with 0.5 mM IPTG. In comparison to the empty pJJW101 vector and uninduced controls, the knockdown plasmids had varied efficiencies at silencing target protein expression when the CRISPRi system was induced (Fig. 2C). Notably, the 162 kDa BbdCas9-myc was visible as a distinct band after Coomassie staining, allowing for the rapid confirmation of system inductions. We also readily observed robust knockdowns of the highly abundant OspA and OspB (31 and 34 kDa, respectively), and to a lesser extent FlaB (37 kDa). Among the nine evaluated sgRNAs, ospAT1 was the only spacer sequence that did not appreciably affect target protein levels. ospAT1’s ineffectiveness is likely multifactorial: First, as observed in other systems, there appears to be an efficiency bias for NT sgRNAs over T sgRNAs, supported by flaBNT2 and ospANT2 sgRNAs leading to the strongest reductions in target protein expression (3, 6). Second, specificity of the 3’ core (PAM-proximal) region of sgRNA spacer sequences has been shown to drive knockdown efficiency. For this reason, the CRISPy-web platform calculates the number of off-target genomic hits containing 0, 1, or 2 bp mismatches for the 13 bp core of each potential 20 bp spacer sequence in a selected ORF (Table 2). Third, a GC core content ≥30% leads to more efficient sgRNA annealing to the target sequences (38–41). ospAT1 had the least favorable values of the tested sgRNAs in all three categories.

**Table 1.**
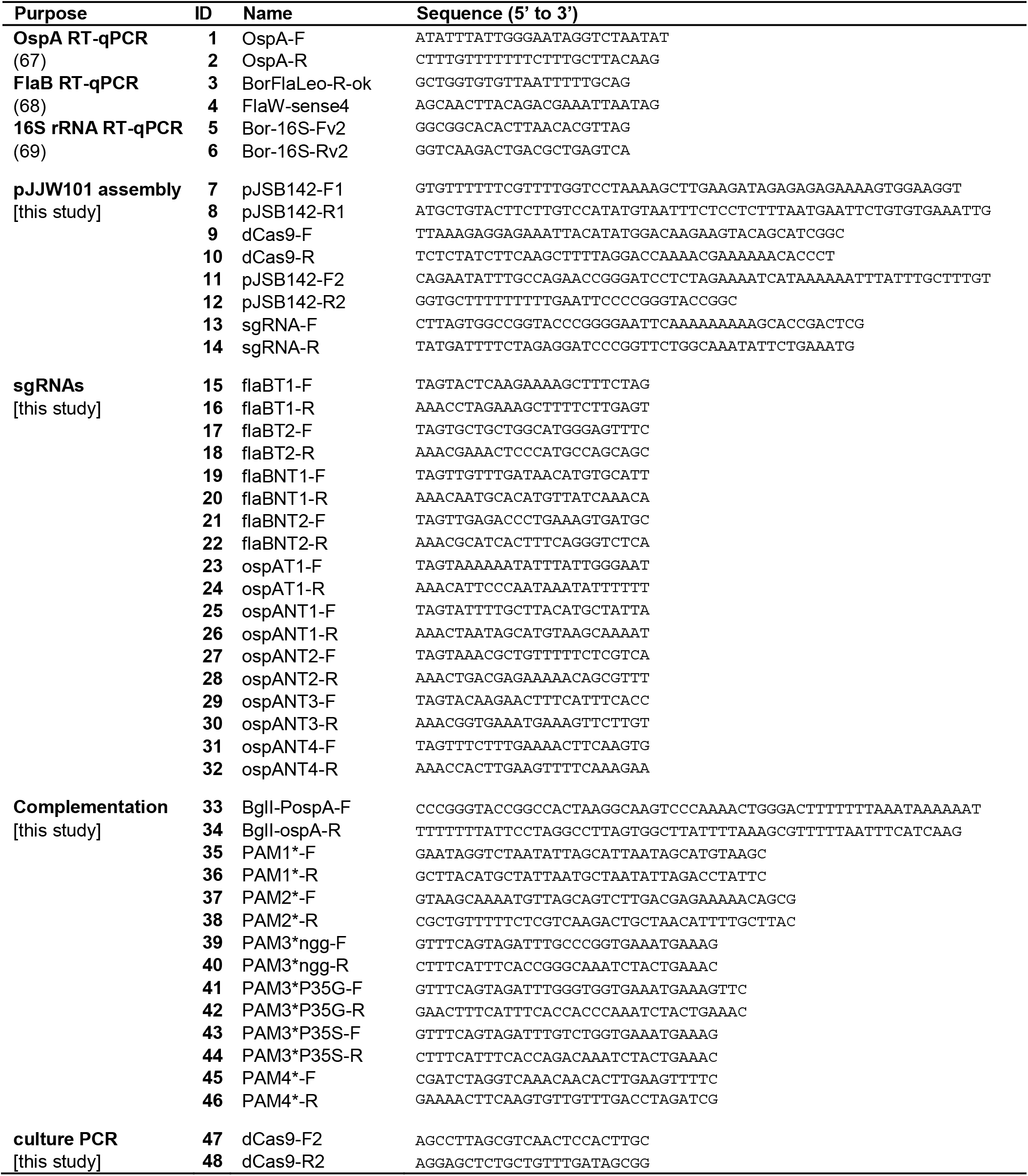
Oligonucleotides used in this study.

**Table 2.**
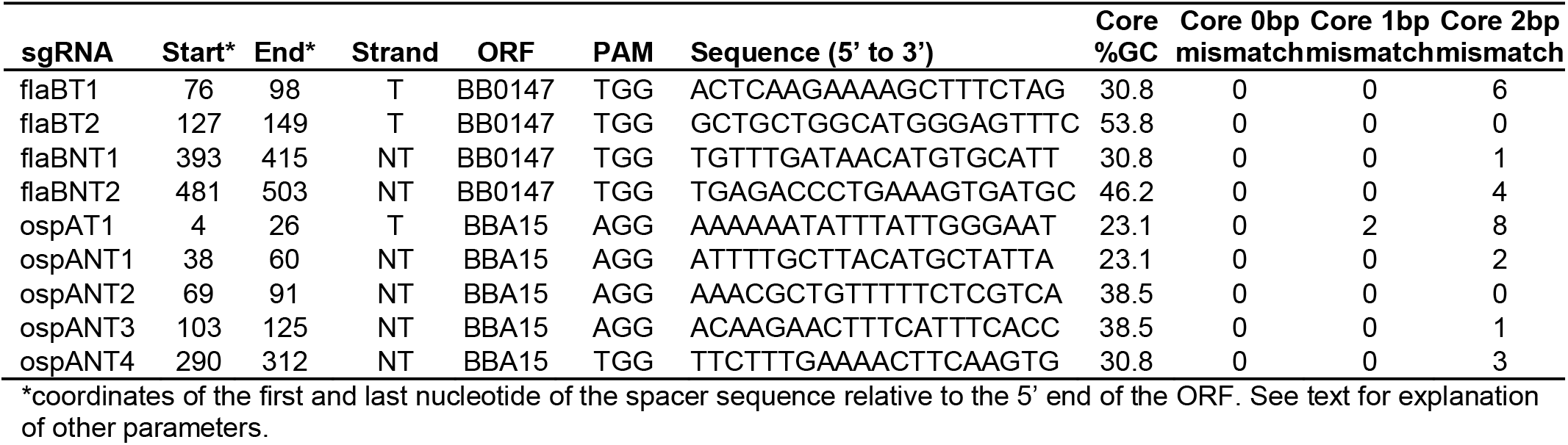
sgRNA spacer sequences used in this study.

**Table 3.** PAM site mutations.

| sgRNA | sgRNA-matching PAM site | PAM* name | PAM* site† | amino acid change |
| --- | --- | --- | --- | --- |
| ospANT1 | AGG | PAM1* | ATG | None |
| ospANT2 | AGG | PAM2* | AGA | None |
| ospANT3 | AGG | PAM3a* | GGG | None |
|  |  | PAM3b* | ACC | Pro to Gly |
|  |  | PAM3c* | AGA | Pro to Ser |
| ospANT4 | TGG | PAM4* | TGT | None |

†Mismatching nucleotides are underlined

**Table 4.** Plasmids used in this study.

| ID | Plasmid | sgRNA | Complementing allele | Knockdown |
| --- | --- | --- | --- | --- |
| 1 | pJJW101 | None | None | None |
| 2 | pJJW101-flaBT1 | flaBT1 | None | FlaB |
| 3 | pJJW101-flaBT2 | flaBT2 | None | FlaB |
| 4 | pJJW101-flaBNT1 | flaBNT1 | None | FlaB |
| 5 | pJJW101-flaBNT2 | flaBNT2 | None | FlaB |
| 6 | pJJW101-ospAT1 | ospAT1 | None | OspAB |
| 7 | pJJW101-ospANT1 | ospANT1 | None | OspAB |
| 8 | pJJW101-ospANT2 | ospANT2 | None | OspAB |
| 9 | pJJW101-ospANT3 | ospANT3 | None | OspAB |
| 10 | pJJW101-ospANT4 | ospANT4 | None | OspAB |
| 11 | pASP151 | ospANT1 | WT *ospA* | OspAB |
| 12 | pASP152 | ospANT2 | WT *ospA* | OspAB |
| 13 | pASP153 | None | PAM1* *ospA* | None |
| 14 | pASP154 | None | PAM2* *ospA* | None |
| 15 | pASP155 | None | WT *ospA* | None |
| 16 | pASP156 | ospANT1 | PAM1* *ospA* | OspB only |
| 17 | pASP157 | ospANT2 | PAM1* *ospA* | OspAB |
| 18 | pASP158 | ospANT1 | PAM2* *ospA* | OspAB |
| 19 | pASP159 | ospANT2 | PAM2* *ospA* | OspB only |
| 20 | pBTM151 | ospANT3 | WT *ospA* | OspAB |
| 21 | pBTM152 | ospANT4 | WT *ospA* | OspAB |
| 22 | pBTM153 | ospANT3 | PAM3a* *ospA* | OspAB |
| 23 | pBTM153e1 | ospANT3 | PAM3b* *ospA* | OspB only |
| 24 | pBTM153e2 | ospANT3 | PAM3c* *ospA* | OspB only |
| 25 | pBTM154 | ospANT4 | PAM4* *ospA* | OspB only |
| 26 | pBTM155 | ospANT3 | PAM2* *ospA* | OspAB |
| 27 | pBTM156 | ospANT4 | PAM2* *ospA* | OspAB |

**Figure 2.**
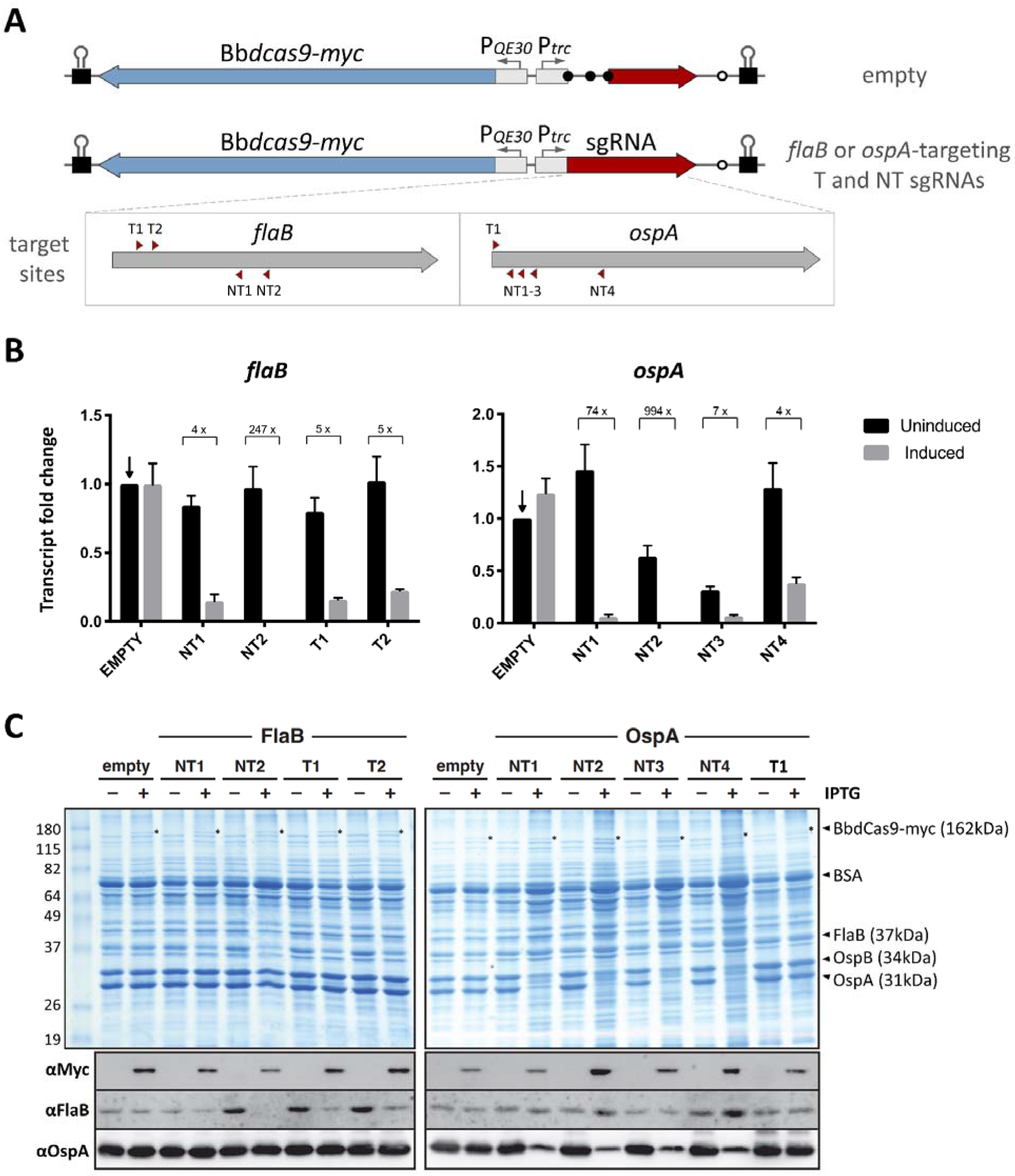
CRISPRi knockdowns of *flaB* and *ospA*. **(A)** Schematic of the pJJW101 knockdown functional region as an empty vector control or with template (T) & non-template (NT) strand gene-targeting sgRNAs. The target sites for each sgRNA (red triangles) are shown proportional to their respective ORF. **(B)** *Borrelia* cells carrying CRISPRi system plasmids having either no gene-targeting sgRNA (empty) or one of eight unique gene-targeting sgRNAs (against *flaB* or *ospA*) were induced with 0.5mM IPTG for 36-hours in comparison to uninduced controls. Quantification of *flaB, ospA*, and 16S rRNA transcripts was done by RT-qPCR. Targeted *flaB* and *ospA* transcripts were normalized against the 16S rRNA endogenous control. Data are expressed as the average fold change of targeted genes by different sgRNAs relative to the amount of transcript present in uninduced empty samples (indicated by arrows). Error bars show the mean + SEM for three independent experiments. **(C)** Coomassie-stained SDS-PAGE gels and Western immunoblots of whole cell lysates prepared from pJJW101- transformed *Borrelia* cells incubated with (+) or without (-) 0.5mM IPTG for 36 hours.

We next harvested total RNA from spirochetes with CRISPRi systems that resulted in strong protein silencing to more directly analyze their knockdown efficiencies at the transcript level by RT-qPCR (Fig. 2B). While generally in line with the immunoblot data (Fig. 2C), the analysis showed 4-fold, 7-fold, and 74-fold decreases in *ospA* transcript abundance for ospANT4, ospANT3, and ospANT1 inductions, respectively. This supports the preference of effective sgRNAs for promoter-proximal targets, but it also indicates that even relatively low transcriptional silencing can be sufficient to produce a phenotype at the protein level. Conversely, the visually greater protein knockdown efficiency of flaBT1 compared to flaBNT1 was measured as only a 25% greater transcript knockdown. These inconsistencies may originate from unique post-transcriptional and post-translational regulation mechanisms. As shown for the *ospAB* operon, variations of *ospAB* mRNA levels among strains or even within cultures of the same strain can still have equivalent protein levels, likely as a result of the same regulatory mechanisms (42–44). Thus, overall CRISPRi sgRNA efficiency should be adjudicated on phenotypic readouts at the protein level whenever possible.

At this point, our CRISPRi platform allowed for the controlled repression of *B. burgdorferi* genes. Yet, as the polar effects of *ospA*-targeting sgRNAs on OspB expression showed, a strict fulfillment of Koch’s molecular postulates in this system was limited to monocistronic genes or the last gene in an operon. Thus, we explored whether we could augment the platform to allow for specific *in trans* complementation. Using the *ospAB* operon for proof-of-principle experiments, we took advantage of the strict protospacer adjacent motif (PAM) requirement for sgRNA-mediated gene targeting of dCas9 (45). The pJJW101 empty vector and its four ospANT sgRNA derivatives were modified by directional insertion of different *ospA* variants under their own promoter (*P_ospA_-ospA*) into a BglI site downstream of the sgRNA module (Fig. 3A). The *P_ospA_-ospA* copy PAM sites for each of the four ospANT sgRNAs were mutated individually, leading to *ospA* PAM* alleles that were expected to be resistant to targeting by their cognate sgRNAs (Table 3). The resulting series of recombinant pJJW101 derivatives (Table 4, plasmid IDs 13-27) contained both matched (e.g., ospANT2 and *ospA* PAM2*) and mismatched (e.g., ospANT2 and *ospA* PAM1*) combinations of sgRNA and complementing *ospA* alleles to evaluate specificity of the complementation system. *B. burgdorferi* clones harboring the plasmids were cultured for 48 hours with or without CRISPRi system induction, and their protein profiles were assessed by SDS-PAGE and immunoblotting (Figs. 3B and 3C). As expected, sgRNA-lacking (empty) control constructs with either wild-type (WT), PAM1* or PAM2* *ospA* variants failed to block OspA and OspB expression. The observed increase in OspB levels upon complementation with the PAM2* *ospA* variant remains enigmatic but could be explained by a previously described OspA-independent OspB expression mechanism (46). Next, we compared the ability of matched and mismatched sgRNA/PAM* *ospA* combinations to selectively restore OspA expression, with WT *ospA* allele constructs used as internal negative controls. As expected, OspA expression was only restored in matched constructs while WT *ospA-*complemented and mismatched constructs retained a dual OspAB knockdown phenotype, notably despite the plasmid-derived increase in *ospA* copy number. While the PAM1*, PAM2*, and PAM4* *ospA* mutations were synonymous, the PAM3 site overlapped with a proline codon, eliminating the possibility for a synonymous change. We therefore generated a series of three mutants: PAM3a* represented a synonymous PAM site “wobble” (NGG to NGG) mutation, which failed to complement OspA expression in an ospANT3-mediated knockdown. The other two nonsynonymous PAM3b* (Pro35Gly) and PAM3c* (Pro35Ser) *ospA* alleles complemented the ospANT3 knockdowns with fully mutant OspA proteins upon system induction. Notably, PAM3c* *ospA* is identical to a naturally occurring OspA homolog from *B. burgdorferi* strain CA7 (GenBank AAA20957.1) That OspA variant is predicted by Alphafold (47) to fold almost identically to *B. burgdorferi* B31 OspA (Fig. 3D) with a slight angular offset that is exacerbated for in the PAM3b* (Pro35Gly) variant (Supplementary file 3). This set of experiments fully validated the *B. burgdorferi* CRISPRi PAM* system for gene silencing and selective gene complementation.

**Figure 3.**
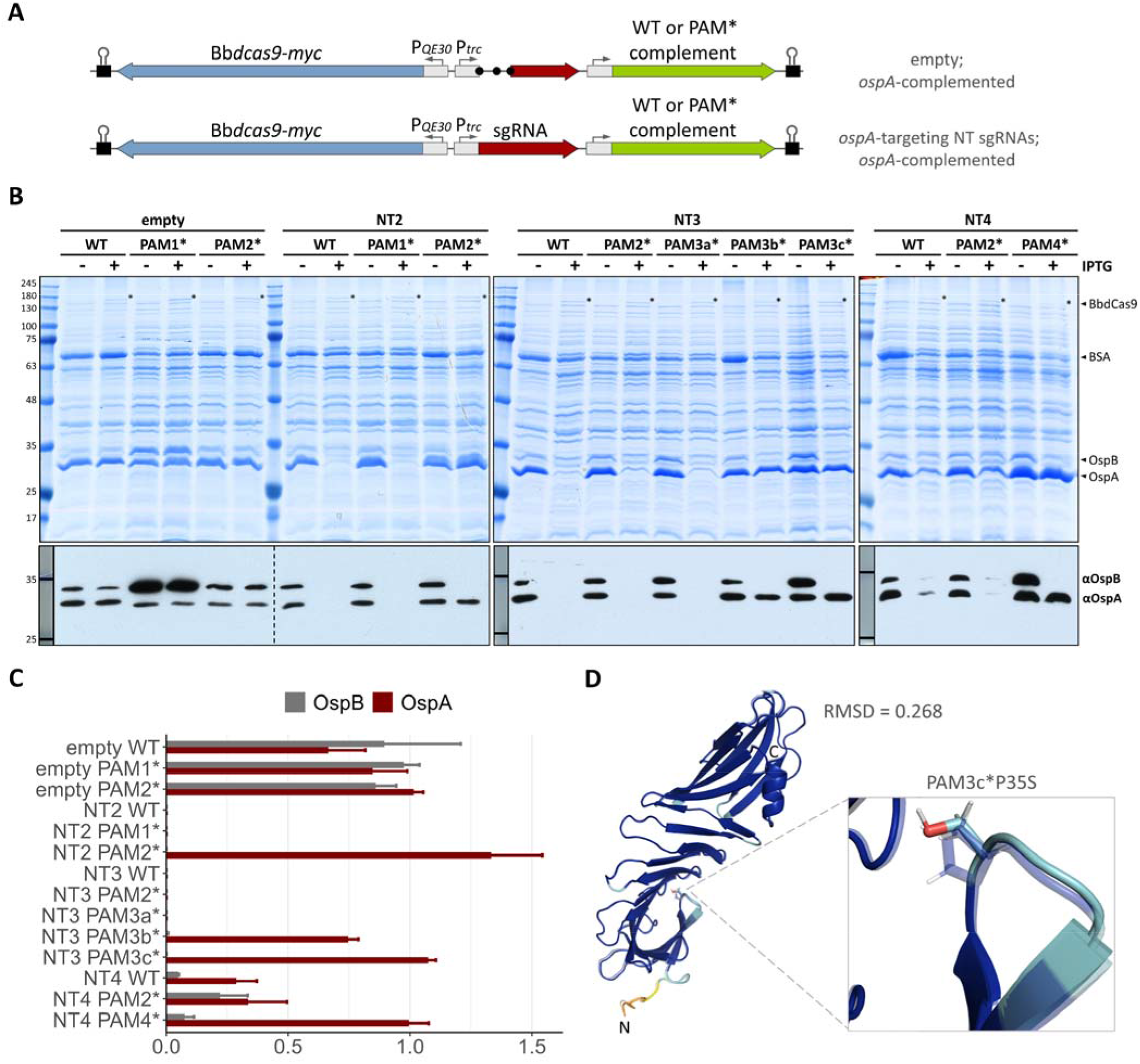
CRISPRi knockdowns of the *ospAB* operon with PAM* *ospA* allele-mediated complementation. **(A)** Schematic of the complemented pJJW101 knockdown functional region as an empty vector control or with gene-targeting sgRNAs. **(B)** *Borrelia* cells carrying either wild type (WT) or PAM mutant (PAM*) *ospA-*complemented CRISPRi system plasmids having either no gene-targeting sgRNA (empty) or one of three unique *ospA*-targeting sgRNAs (NT2, NT3, or NT4) were induced (+) with 0.5 mM IPTG for 48-hours in comparison to uninduced (-) controls. Whole cell lysates were analyzed by Coomassie-stained SDS-PAGE gels and anti-OspA and anti-OspB immunoblots. **(C)** Immunoblots for the *ospA-*complemented CRISPRi system inductions (including data in Fig. 3B) were quantified with ImageJ to visualize knockdown proportions as the ration of induced/uninduced protein levels. Error bars show the mean ± SEM for three independent experiments. **(D)** AlphaFold prediction for mature P35S OspA mutant is superimposed onto the prediction for mature WT OspA (transparent) and colored by predicted local difference distance test (pLDDT) score for each residue. PyMOL calculated RMSD for the superimposition is shown.

## DISCUSSION

The CRISPRi PAM* system described here further augments the genetic toolbox for the spirochetal pathogen *B. burgdorferi*, providing a method for inducible knockdowns with optional *in trans* complementation. All necessary factors for knockdown and complementation are encoded on a single recombinant plasmid, making the system convenient for any *B. burgdorferi* strain background and–with some minor adaptations–transferable to genetically amenable relapsing fever *Borrelia* (15, 48–51).

A recent study by Takacs *et al*. (10) also observed toxicity of the original *S. pyogenes* dCas9 gene in *B. burgdorferi*. While that study used weakened promoters to reduce SpydCas9 expression to tolerated levels, we were able to overcome heterologous protein toxicity by codon optimization of the GC-rich SpydCas9 gene (56% G+C) to match the AT-rich *B. burgdorferi* genome (Supplementary file 1). This suggests that SpydCas9’s toxicity in *B. burgdorferi* is due to rare codon usage (35) which is likely exacerbated by the large size of the transcript/protein. Codon-optimized BbdCas9 allowed us to maintain use of the original *P_QE30_* (T5/*lac*) hybrid promoter for IPTG-driven BbdCas9 expression. As a result, BbdCas9 protein was clearly detectable in Coomassie-stained SDS-PAGE gels of induced whole cell protein samples, but it remained undetectable by immunoblot (via a C-terminal C-myc epitope tag) under non-inducing conditions (Fig. 2C), indicating tightly controlled expression. Our system also added an additional level of control by putting the sgRNA module under a *trc* (*trp/lac*) hybrid promoter, which ensures that both dCas9 and sgRNA CRISPRi system components remain repressed until being simultaneously induced by IPTG. Additional modifications that replace the currently *lac*-based promoters with a *tet*-responsive hybrid promoter (26) might render the system more titratable to match the CRISPRi systems for some other bacteria (6, 34).

The CRISPy-web interface provided an accessible and relatively simple tool for identifying potential sgRNA spacers. When asked to query a genomic sequence like the artificial concatenated *B. burgdorferi* strain B31 contig used here, it produces a ranked list of spacers for each selected ORF, using the lowest number of genomic off-target core hits as the most important parameter for selection and ranking (37). Our set of experiments continue to support the general rules that more efficient spacers also (i) target the non-template strand, (ii) bind closer to the targeted gene’s promoter region, and (iii) have ≥30% GC content within their core region.

As demonstrated in this study, the *B. burgdorferi* CRISPRi PAM* approach can be used to restore expression of a targeted gene. Mutation of the PAM sequence alone is sufficient to generate a “CRISPRi-resistant” complementing allele. However, it should be noted that NT or T sgRNA PAM sites overlapping Pro or Gly codons, respectively, will require nonsynonymous mutations to produce PAM* sites. In these cases, structural information and comparative genomics can be used to guide sgRNA selection and PAM* mutant design. For example, in our ospANT3 PAM* complementation series, the affected OspA Pro35 lies in a loop connecting two β-sheets (Fig. 3D and Supplementary file 3) where substitution mutations are less likely to disrupt structure and destabilize the protein. We first used a BLAST search to identify a naturally occurring OspA Pro35Ser variant in *Borrelia burgdorferi* strain CA7 (52), which informed the design of the PAM3c* *ospA* allele (Table 3). The PAM3b* allele replaced Pro35 with the small and flexible Gly residue. As predicted, both complementing OspA mutants were readily detected by SDS-PAGE and Western immunoblotting (Fig. 3B). Alternatively, CRISPRi resistant alleles can be generated by disrupting complementarity to the core spacer sequence. A machine learning approach for spacer sequence mismatching has been used to determine which substitutions will most likely reduce sgRNA activity in *E. coli* and *Bacillus subtilis* (53), and a combination of PAM and spacer mutations was recently used in a screen for *Mycobacterium tuberculosis* antimicrobial resistance mechanisms (54).

One obvious application of the *B. burgdorferi* CRISPRi PAM* system is to ensure that a CRISPRi-mediated phenotype is indeed due to silencing of the targeted gene alone. This will be particularly important when assessing potential polar effects in phenotypical studies of genes that, based on RNASeq data (55), are either not at the end of a polycistronic transcript or not independently transcribed. If complementation with the PAM* allele does not restore the WT phenotype, the same pJJW101-based CRISPRi system can be used to complement with the operon’s non-targeted genes. The currently largest pJJW101-related recombinant plasmid is about 14 kb in size (22), which means that pJJW101 should accommodate at least 3 kb of complementing sequence. Larger operons might require placement of the system’s BbdCas9 and BbLacI expression cassettes on stable *B. burgdorferi* replicons as in the study by Takacs et al. (10).

Most intriguingly, the *B. burgdorferi* CRISPRi PAM* system can be employed to simultaneously silence a gene and complement it with a mutant allele, similar to the above-described replacement of strain B31 OspA by a mutant variant from strain CA7. This system application will be particularly helpful in structure-function studies of essential *B. burgdorferi* proteins. Traditionally, construction of an essential gene knockdown strain first required adding an inducible target gene allele on a recombinant plasmid, resulting in a merodiploid intermediate. This intermediate then had to be mutagenized further to disrupt the endogenous target gene locus by allelic exchange with an antibiotic cassette, finally yielding the haploid conditional knockout. In structure-function studies, this two-step protocol had to be repeated for each tested mutant. CRISPRi PAM* reduces this protocol to a single transformation with a single plasmid. This increased efficiency in *B. burgdorferi* molecular genetics also opens the door to generating CRISPRi PAM* plasmid-based mutant libraries in a Mut-Seq approach (56) to probe essential residues within essential proteins. To that end, we are currently using CRISPRi PAM* for structure-function studies of essential protein secretion and envelope homeostasis pathway components in *B. burgdorferi*.

## MATERIALS & METHODS

### Bacterial strains, growth conditions, and recombinant plasmid purification

Chemically competent NEB5-alpha cells (NEB, C2987H) were transformed with recombinant plasmids per manufacturer instructions and grown at 37°C on selective LB agar plates (BD, 244520) and in LB broth (Fisher, BP1426) containing 30μg/mL kanamycin (Sigma, K4000). Plasmid DNA was isolated from *E. coli* clones using a Miniprep kit (Macherey-Nagel, 740588) and verified by single pass primer extension sequencing (ACGT Inc.) with either generic or custom DNA oligonucleotide primers (Integrated DNA Technologies). Verified plasmids were purified using a Midiprep kit (Macherey-Nagel, 740412) and concentrated by standard sodium acetate/ethanol precipitation (REF Sambrook) to ≥500ng/μL for transformation into the *B. burgdorferi* strain type strain clone B31-e2 (57). *B. burgdorferi* was grown in liquid sterile filtered Barbour-Stonner-Kelly-II (BSK-II) medium (58, 59) containing 9.7 g/L CMRL-1066 (US Biological, C5900-01), 5.0 g/L neopeptone (Gibco, 211681), 6.6 g/L HEPES sodium salt (Fisher, BP410), 0.7 g/L citric acid (Sigma, C-8532), 5.0 g/L dextrose anhydrous (Fisher, BP350), 2.0 g/L yeastolate (Gibco, 255772), 2.2 g/L sodium bicarbonate (Fisher, BP328), 0.8 g/L sodium pyruvate (Fisher, AC132155000), 0.4 g/L *N*-acetylglucosamine (Sigma, A3286), 25 mg/L phenol red (Sigma, P-3532), and 50.0 g/L bovine serum albumin (Gemini, 700-104P) at pH 7.6-7.7, with 60□mL/L heat-inactivated rabbit serum (Pel Freez, 31126), and 200 mL/L 7% gelatin (Gibco, 214340) added before use. The cultures were incubated at 34°C under 5% CO_2_ in a humidified incubator. Recombinant B31-e2 clones were propagated in BSK-II with 200μg/mL kanamycin for plasmid maintenance.

### Transformation and clonal selection of *B. burgdorferi*

Electrocompetent B31-e2 cells were transformed by electroporation (60) with 1-5μg of plasmid in 2mm gap cuvettes (Thermo, 5520) using a BioRad MicroPulser on EC2 setting, consistently measuring 2.49 kV/cm field strength and 5.20-5.60 ms pulse times. Electroporated cells were immediately resuspended in 12mL of pre-warmed BSK-II and allowed to recover at 34°C for 18-20 hrs. Clonal selection of transformants was carried out by adding the 12 mL recovered culture to 35mL BSK-II with a final concentration of 200μg/mL kanamycin, followed by plating into 96-well microtiter plates and 8-16 days of incubation (61). Culture-positive wells were expanded into 6 mL of selective BSK-II and allowed to reach stationary phase for verification of plasmid acquisition by direct PCR of cultured *B. burgdorferi* cells (62) using QuickLoad 2x Taq Master Mix (NEB, M0271) and primers 49 and 50 (Table 1). Positive clones were flash frozen on dry ice in BSK-II containing 10% DMSO (Sigma, D2438) and stored at -80°C.

### CRISPRi system plasmid construction

pJJW101 was constructed using two stepwise NEBuilder HiFi assemblies (NEB, E5520S) followed by restriction and ligation. All used oligonucleotide primers are referred to by their ID number listed in Table 1. For the first assembly, the vector backbone of pJSB142, a kanamycin resistance-conferring derivative of pJSB104 (25), was amplified using primers 7 and 8 and combined with an SpydCas9 ORF module amplified by primers 9 and 10 from pTn7C107 (34). For the second assembly, the resulting pJSB142-dCas9 shuttle vector plasmid was amplified in its entirety with primers 11 and 12 to combine with a P_*trc*_-driven sgRNA module amplified by primers 13 and 14 from pTn7C107, resulting in pJJW100. Next, a standard format codon usage table for *B. burgdorferi* B31 was obtained from the codon usage database (63) for species-specific codon optimization of dCas9 using the random guided codons parameter on OPTIMIZER (Supplementary material 1) (36). This codon-optimized version of *dcas9* (Bb*dcas9*) was synthesized and cloned in pBluescript II KS (+) with flanking 5’ *NdeI* and 3’ *NotI* restriction endonuclease sites (GenScript), yielding pBS-Bbdcas9. Finally, the *NdeI-NotI* SpydCas9 fragment of pJJW100 was replaced by the *NdeI-NotI* Bb*dcas9* fragment from pBS-Bbdcas9, yielding pJJW101.

### sgRNA spacer sequence design and insertion into pJJW101

The 20-nucleotide spacer sequences for our sgRNAs were designed using the CRISPy-web interface on an artificially concatenated sequence of the entire *Borrelia burgdorferi* B31 genome having individual genetic elements separated by “NNNNNN” (Supplementary file 2) (37). Four sgRNAs, two each complementary to the non-template (NT) and template (T) strands, were chosen for targeting *flaB*. One T and four NT sgRNAs were chosen for targeting *ospA* (Table 2). The 5’ to 3’ spacer sequences listed in the CRISPy-web output were used to design oligonucleotide pairs: Forward oligos added 5’-TAGT overhangs to the spacer sequence, while reverse oligos added 5’-AAAC overhangs to the reverse complement of the spacer sequence (Table 1, oligo IDs 15-32). Complementary pairs of single-stranded oligonucleotides were mixed at a final concentration of 2.4μM, denatured at 95°C for 5 min, and annealed by gradual cooling to room temperature (https://portals.broadinstitute.org/gpp/public/resources/protocols). The resulting double-stranded oligos had 5’ extensions that were complementary to the non-palindromic 3’ overhangs generated by BsaI-HFv2 (NEB, R3733) digestion of pJJW101, facilitating efficient directional ligation of hybridized sgRNA primers and cut pJJW101 vector.

### WT and PAM* *ospA* allele generation

The coding region for OspA with its native intergenic promotor was amplified by PCR with Q5 thermostable high-fidelity polymerase (NEB, M0491) from B31-e2 genomic DNA with primers 35 and 36 (Table 1) containing 5’-BglI overhangs (Table 1), and the resulting amplicon was cloned into pCR-Blunt II-TOPO (Invitrogen, 451245). This plasmid was then used as template for Splicing by Overlap extension (SOE) PCR with Q5 polymerase, flanking primers 35 and 36, and pairs of mutagenic complementary PAM* primers (Table 1, primers 37 to 48) to generate six PAM* versions of the BglI-flanked *P_ospA_-ospA* amplicon. Synonymous mutations were introduced into the PAM sites wherever possible. E.g., the PAM1* site 5’-AGG-3’ to 5’-ATG-3’ corresponded to a synonymous 5’-GCC-3’ to 5’-GCA-3’ Ala codon change on the complementary strand. PAM3* could not be made synonymously, an issue for non-template sgRNAs that occurs when the reverse complement 5’-NGG-3’ PAM site aligns with a 5’-CCN-3’ Pro codon. To overcome this, the *ospA* (ORF BBA15) nucleotide sequence was used in an NCBI blastx query with default search parameters and a filter for *Borrelia* sequences. Sifting through the alignments tab with alignment view set to “Query-anchored with dots for identities” allowed for quick identification of a naturally occurring P35S allelic variant in *Borrelia burgdorferi* strain CA7 which was used for the PAM3c* *ospA* allele.

### Complementation of CRISPRi constructs with WT and mutant PAM* alleles

pJJW101 derivatives containing one of the four NT *ospA*-targeting sgRNAs were cut with BglI (NEB, R0143), dephosphorylated with Quick-CIP (NEB, M0525), and ligated with T4 ligase (NEB, M0202) at a 1:2 ratio with either WT or PAM* BglI-cut *PospA-ospA* amplicons. Isolated plasmid DNA from *E. coli* transformants was checked for insert by digestion with BglI prior to confirmatory sequencing with primer 14 (Table 1).

### CRISPRi system inductions and growth curves

Recombinant *B. burgdorferi* strain stocks grown in selective BSK-II were diluted in selective medium with or without 0.5 mM IPTG and harvested after 36 or 48 hours of incubation. Whole cell lysates were checked for overall changes in transcript and protein levels by SDS-PAGE and Western immunoblotting as detailed below. Cell density was assessed by direct counting of bacterial cultures diluted 10- to 100-fold in PBS in a Petroff Hauser counting chamber under phase contrast microscope (Nikon Eclipse E400).

### RT-qPCR for transcript expression analysis

Total RNA was extracted from *B. burgdorferi* cells with TRIzol Reagent (Invitrogen, 15596026) according to the manufacturer’s instructions after a 30-minute 3,000 x *g* swinging-bucket centrifugation at room temperature. Residual DNA contamination was removed by a 1-hour DNase I treatment (Thermo Scientific, 18047019) followed by phenol-chloroform extraction (Ambion, AM9720) and standard ammonium acetate/ethanol precipitation (64). RNA samples were used for reverse transcription and quantification of *ospA, flaB*, and 16s rRNA transcripts by the Luna Universal One-Step RT-qPCR kit (NEB, E3005), according to the manufacturer’s instructions on an Applied Biosystems 7500 Fast Real-Time PCR System. The primers to amplify *flaB, ospA* and 16S RNA transcripts are shown and referenced in Table 1 (primers 1-6). Transcript levels were validated and normalized against 16S rRNA transcript with fold changes calculated using the comparative CT (2^-ΔΔCT^) method for quantification.

### SDS-PAGE and Immunoblotting for protein expression analysis

*B. burgdorferi* cells grown to late logarithmic or early stationary phase were harvested by centrifugation in a swinging bucket at 3,000 x *g* for 30 min at room temperature. Harvested cells were washed twice with dPBS + 5mM MgSO4 to remove culture medium BSA and resuspended in standard 1 x SDS sample buffer (64). Whole cell lysates were resolved by 12% sodium dodecyl sulfate polyacrylamide gel electrophoresis (SDS-PAGE) and visualized by Coomassie staining (Fisher, BP3620-1). For immunoblotting, proteins were transferred electrophoretically to nitrocellulose blotting membrane (GE, 10600002) using a BioRad Mini Trans-Blot apparatus at 100V for 1 hour with pre-chilled transfer buffer (25mM Tris, 200mM Glycine, 20% methanol). Membranes were blocked for 30 min in TBST buffer (25 mM Tris, 150 mM NaCl, 0.05% Tween 20, pH 7.2) with 5% dry milk before incubating on a rocker overnight at 4°C with rat anti-FlaB polyclonal antiserum (1:4,000 dilution; a gift from M. Caimano, University of Connecticut Health Center), mouse anti-c-Myc 9E10 (1:1,000 dilution; Fisher, MA1-980), mouse anti-OspA (1:10,000 dilution, H5332 (65)), or mouse anti-OspB (1:5,000 dilution, H6831 (66)) primary antibodies. After three 10-minute washes with TBST, the blots were incubated at room temperature on a rocker for 1 hr with secondary anti-mouse IgG-HRP (Sigma, A4416) or anti-rat IgG-HRP (Thermo, 31470) antibody. three additional 10-minute TBST washes, membranes were allowed to react with Super Signal West Femto (Thermo, 34096) substrate per manufacturer instructions and the signal was detected by exposure to autoradiography film (Midsci, XC59X).

**Figure 4.**
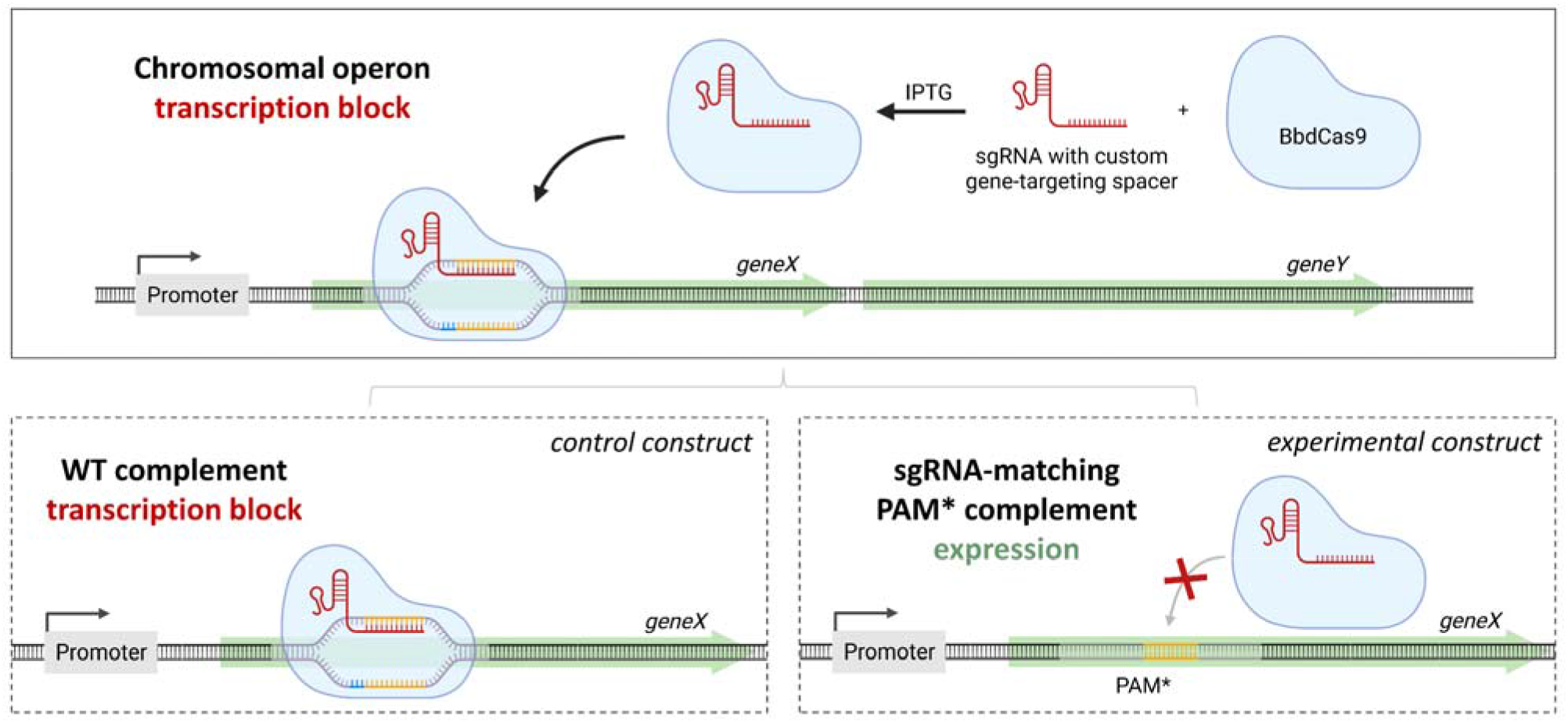
CRISPRi PAM* system overview. After IPTG-induced expression, the BbdCas9-sgRNA complex locates complementary protospacer sequences (gold) with 5’-NGG PAM sites (blue) to sterically block transcription elongation. When the CRISPRi system is complemented with the wild type (WT) allele of the targeted gene, the complementing allele is subject to the same transcription blockade as the genomic target (control construct). However, when the system is complemented with a PAM* mutant allele of the targeted gene, the BbdCas9-sgRNA complex can no longer bind to that allele, and PAM* allele transcription is permitted (experimental construct) while the transcription blockade of the genomic target is maintained. Created with BioRender.com.

## ACKNOWLEDGMENTS

We thank Jon Blevins (University of Arkansas for Medical Sciences, Department of Microbiology & Immunology, Little Rock, Arkansas) for recombinant plasmid constructs. WRZ and BS conceived and designed the study; JMP provided materials; BM, JJW, HH, and ASP acquired the data; BM, HH and WRZ analyzed and interpreted the data; BM and WRZ wrote the manuscript with contributions from BS and JMP. This work was supported by NIH/NIAID grant R21AI144624 and a KUMC Lied Basic Science Pilot Grant to WRZ and NIH/NIAID grants R01AI144126, 3R01AI144126-03S1, and R21AI147139 to BS.

