## Supplemental files 1 and 3 for "Inducible CRISPRi-based operon silencing and selective *in trans* gene complementation in *Borrelia burgdorferi*"

**SUPPLEMENTARY DATA**

**Supplementary File 1**. **OPTIMIZER output table for BbdCas9.**

| **Type** | **Sequences** | **CAI** | **ENc** | **%GC** | **%AT** |
| --- | --- | --- | --- | --- | --- |
| Query | MDKKYSIGLAIGTNSVGWAVITDEYKVPSKKFKVLGNTDRHSIKKNLIGALLFDSGETAEATRLKRTARRRYTRRKNRICYLQEIFSNEMAKVDDSFFHRLEESFLVEEDKKHERHPIFGNIVDEVAYHEKYPTIYHLRKKLVDSTDKADLRLIYLALAHMIKFRGHFLIEGDLNPDNSDVDKLFIQLVQTYNQLFEENPINASGVDAKAILSARLSKSRRLENLIAQLPGEKKNGLFGNLIALSLGLTPNFKSNFDLAEDAKLQLSKDTYDDDLDNLLAQIGDQYADLFLAAKNLSDAILLSDILRVNTEITKAPLSASMIKRYDEHHQDLTLLKALVRQQLPEKYKEIFFDQSKNGYAGYIDGGASQEEFYKFIKPILEKMDGTEELLVKLNREDLLRKQRTFDNGSIPHQIHLGELHAILRRQEDFYPFLKDNREKIEKILTFRIPYYVGPLARGNSRFAWMTRKSEETITPWNFEEVVDKGASAQSFIERMTNFDKNLPNEKVLPKHSLLYEYFTVYNELTKVKYVTEGMRKPAFLSGEQKKAIVDLLFKTNRKVTVKQLKEDYFKKIECFDSVEISGVEDRFNASLGTYHDLLKIIKDKDFLDNEENEDILEDIVLTLTLFEDREMIEERLKTYAHLFDDKVMKQLKRRRYTGWGRLSRKLINGIRDKQSGKTILDFLKSDGFANRNFMQLIHDDSLTFKEDIQKAQVSGQGDSLHEHIANLAGSPAIKKGILQTVKVVDELVKVMGRHKPENIVIEMARENQTTQKGQKNSRERMKRIEEGIKELGSQILKEHPVENTQLQNEKLYLYYLQNGRDMYVDQELDINRLSDYDVDAIVPQSFLKDDSIDNKVLTRSDKNRGKSDNVPSEEVVKKMKNYWRQLLNAKLITQRKFDNLTKAERGGLSELDKAGFIKRQLVETRQITKHVAQILDSRMNTKYDENDKLIREVKVITLKSKLVSDFRKDFQFYKVREINNYHHAHDAYLNAVVGTALIKKYPKLESEFVYGDYKVYDVRKMIAKSEQEIGKATAKYFFYSNIMNFFKTEITLANGEIRKRPLIETNGETGEIVWDKGRDFATVRKVLSMPQVNIVKKTEVQTGGFSKESILPKRNSDKLIARKKDWDPKKYGGFDSPTVAYSVLVVAKVEKGKSKKLKSVKELLGITIMERSSFEKNPIDFLEAKGYKEVKKDLIIKLPKYSLFELENGRKRMLASAGELQKGNELALPSKYVNFLYLASHYEKLKGSPEDNEQKQLFVEQHKHYLDEIIEQISEFSKRVILADANLDKVLSAYNKHRDKPIREQAENIIHLFTLTNLGAPAAFKYFDTTIDRKRYTSTKEVLDATLIHQSITGLYETRIDLSQLGGD | - | - | - | - |
| Optimized | ATGGATAAAAAATACTCAATTGGTTTAGCTATAGGAACCAATAGCGTTGGTTGGGCAGTTATTACTGATGAGTATAAAGTTCCATCTAAAAAATTTAAAGTTCTCGGAAACACTGATAGACATTCCATAAAAAAAAACTTAATAGGAGCTCTGCTGTTTGATAGCGGTGAAACTGCAGAGGCAACAAGATTGAAGAGAACTGCAAGAAGGCGTTATACTAGGAGAAAGAATAGAATATGTTATCTTCAAGAAATATTTAGTAATGAAATGGCTAAAGTTGATGATTCTTTCTTTCACAGACTTGAGGAAAGCTTCCTTGTAGAAGAAGATAAGAAACATGAAAGACACCCCATTTTTGGAAATATTGTAGATGAAGTTGCATATCATGAAAAATATCCAACTATATATCATTTGAGAAAAAAGTTAGTTGACTCAACTGATAAAGCTGACCTTAGGTTAATATATCTTGCGTTAGCACATATGATTAAGTTTAGGGGACATTTCCTAATTGAAGGAGATTTAAATCCTGATAATTCTGACGTTGACAAATTGTTTATTCAACTTGTACAGACATATAATCAATTGTTTGAGGAAAATCCGATTAATGCAAGTGGAGTTGACGCTAAGGCTATTTTATCTGCAAGACTTTCTAAATCCAGGAGATTAGAAAACTTGATTGCTCAACTTCCTGGAGAGAAGAAAAATGGACTTTTTGGAAATCTTATAGCACTAAGCTTAGGATTAACACCGAATTTTAAGAGTAATTTTGACTTAGCTGAGGATGCCAAACTTCAATTGTCTAAAGATACGTATGACGACGATCTTGACAATTTATTAGCTCAAATTGGAGATCAATATGCAGATTTATTTCTTGCAGCTAAAAATTTATCTGATGCCATTTTGTTGTCTGATATTCTAAGAGTAAATACCGAAATAACAAAGGCACCATTGAGTGCAAGTATGATTAAAAGGTATGATGAGCATCATCAAGATTTAACATTGCTTAAGGCTTTAGTAAGACAACAATTACCTGAGAAATACAAAGAAATTTTTTTTGACCAATCAAAAAATGGTTATGCTGGATATATTGATGGTGGTGCTTCACAAGAAGAATTCTATAAATTTATTAAGCCTATCCTAGAAAAAATGGATGGCACAGAAGAATTACTAGTTAAATTGAACAGGGAAGATTTATTAAGAAAACAAAGAACATTTGATAACGGTAGTATTCCGCATCAAATTCACTTGGGCGAACTTCATGCTATTCTGAGGAGGCAGGAAGATTTTTATCCCTTCTTAAAGGATAATAGAGAAAAAATTGAAAAAATTTTAACATTTAGGATTCCATATTATGTTGGACCCTTAGCTAGAGGAAATAGTAGATTCGCATGGATGACCCGTAAATCTGAAGAAACTATAACTCCCTGGAATTTTGAAGAAGTAGTTGATAAAGGAGCAAGTGCTCAATCTTTCATAGAAAGAATGACAAATTTTGATAAAAACCTTCCTAATGAGAAAGTTTTACCTAAACATAGTTTATTATATGAGTACTTCACTGTTTATAACGAATTAACAAAAGTGAAATACGTTACAGAGGGTATGAGAAAGCCCGCATTTTTGAGTGGGGAACAAAAAAAAGCGATTGTTGACTTGTTGTTCAAAACAAATCGTAAAGTTACTGTAAAACAGCTAAAAGAAGATTATTTTAAAAAAATTGAGTGCTTTGATTCAGTTGAAATATCTGGAGTTGAAGATCGCTTTAATGCAAGTTTAGGAACTTATCATGACCTTTTAAAAATAATTAAAGATAAAGATTTTTTGGATAATGAAGAAAATGAAGATATATTGGAAGATATAGTTCTGACTCTTACTTTATTTGAAGACAGAGAAATGATAGAGGAGCGTCTTAAAACCTATGCACATCTTTTTGATGATAAGGTTATGAAACAATTAAAAAGGAGGAGATATACTGGGTGGGGAAGACTTTCAAGAAAATTGATAAATGGGATCCGCGATAAACAAAGTGGAAAAACAATTTTGGATTTTCTTAAATCTGATGGCTTTGCCAATCGTAACTTTATGCAATTAATACATGATGACTCACTTACTTTTAAAGAAGATATTCAAAAAGCACAAGTTAGTGGACAAGGTGACAGTCTTCATGAGCATATTGCTAATTTAGCTGGATCTCCTGCTATAAAAAAAGGGATATTGCAAACTGTTAAAGTTGTTGATGAACTCGTTAAAGTAATGGGTAGACATAAACCAGAGAATATTGTCATAGAAATGGCCCGGGAGAATCAGACCACACAAAAGGGACAGAAAAATAGCCGAGAAAGGATGAAACGGATTGAAGAAGGTATTAAAGAACTAGGATCTCAAATTTTGAAAGAACATCCTGTTGAGAATACACAATTACAAAATGAAAAATTGTATCTTTATTATTTACAAAACGGCAGAGATATGTACGTTGATCAAGAGCTTGATATTAACAGATTAAGCGATTACGATGTTGATGCAATAGTGCCTCAATCATTTTTAAAAGATGATAGCATAGATAACAAAGTTCTAACTAGATCAGATAAAAATAGAGGAAAATCAGATAATGTACCATCAGAGGAGGTAGTTAAAAAAATGAAAAACTACTGGAGGCAATTACTGAATGCTAAATTAATAACCCAAAGAAAATTTGATAATTTAACAAAGGCAGAAAGAGGAGGATTGAGTGAACTTGATAAAGCAGGATTTATAAAACGGCAATTAGTTGAGACAAGACAAATAACTAAGCACGTCGCTCAAATTCTTGATAGTAGAATGAATACTAAATATGATGAAAACGATAAATTGATTAGAGAGGTAAAAGTTATTACATTAAAATCAAAATTAGTATCTGATTTTAGAAAAGACTTTCAGTTTTATAAAGTAAGAGAGATTAATAATTATCACCATGCTCATGATGCATATTTAAATGCCGTTGTTGGCACTGCTCTCATAAAAAAATATCCTAAGCTTGAATCGGAATTTGTATATGGAGATTATAAAGTTTATGATGTAAGAAAAATGATTGCCAAATCAGAGCAAGAAATAGGTAAAGCTACTGCAAAATATTTTTTTTATTCTAATATTATGAACTTTTTTAAAACTGAAATTACTTTAGCTAACGGAGAGATAAGAAAAAGACCTTTAATTGAAACTAATGGTGAGACTGGGGAGATAGTTTGGGACAAAGGAAGAGATTTTGCAACTGTTCGAAAAGTATTAAGTATGCCTCAAGTTAATATCGTTAAGAAGACTGAAGTTCAAACAGGTGGATTTTCTAAAGAATCAATCTTACCTAAGAGAAATTCTGATAAACTTATTGCAAGGAAAAAAGATTGGGATCCCAAAAAATATGGCGGGTTTGACAGTCCAACCGTTGCCTATTCAGTATTAGTAGTTGCTAAAGTAGAGAAGGGAAAGTCAAAAAAATTAAAAAGCGTTAAAGAATTGCTTGGAATTACTATTATGGAAAGGTCTAGCTTTGAAAAAAATCCTATTGATTTTTTAGAAGCTAAAGGGTATAAAGAAGTTAAAAAAGATTTAATAATAAAATTACCTAAATACAGCTTGTTCGAGTTGGAAAACGGAAGGAAGAGGATGCTTGCAAGTGCCGGAGAATTACAGAAGGGCAATGAATTAGCACTTCCAAGTAAATATGTAAATTTTTTGTATTTAGCTTCTCATTATGAGAAACTAAAAGGGTCACCTGAAGATAATGAACAGAAACAATTATTTGTTGAACAGCATAAACATTATTTAGATGAAATCATCGAACAAATATCTGAGTTTAGCAAGAGAGTTATACTAGCTGATGCAAATTTGGATAAAGTATTATCAGCTTATAATAAACATAGAGATAAACCAATTAGAGAACAAGCAGAAAATATTATACATTTGTTTACACTTACAAATTTAGGGGCACCAGCTGCTTTCAAATATTTTGACACAACAATTGATAGGAAACGTTATACCTCAACAAAGGAAGTTTTGGATGCAACTCTAATTCATCAGTCTATTACTGGATTATATGAGACCAGAATTGATTTGAGCCAATTAGGGGGAGAT | 0.709 | 44 | 31.8 | 68.2 |

| **Codons** | **Usage** | **Codons** | **Usage** | **Codons** | **Usage** | **Codons** | **Usage** |
| --- | --- | --- | --- | --- | --- | --- | --- |
| GCA (A) | 32 | GCC (A) | 8 | GCG (A) | 2 | GCT (A) | 32 |
| TGC (C) | 1 | TGT (C) | 1 | GAC (D) | 19 | GAT (D) | 79 |
| GAA (E) | 75 | GAG (E) | 33 | TTC (F) | 11 | TTT (F) | 52 |
| GGA (G) | 38 | GGC (G) | 7 | GGG (G) | 10 | GGT (G) | 14 |
| CAC (H) | 5 | CAT (H) | 26 | ATA (I) | 35 | ATC (I) | 6 |
| ATT (I) | 52 | AAA (K) | 123 | AAG (K) | 27 | TTA (L) | 63 |
| TTG (L) | 32 | CTA (L) | 11 | CTC (L) | 3 | CTG (L) | 5 |
| CTT (L) | 34 | ATG (M) | 22 | AAC (N) | 16 | AAT (N) | 54 |
| CCA (P) | 10 | CCC (P) | 6 | CCG (P) | 3 | CCT (P) | 16 |
| CAA (Q) | 42 | CAG (Q) | 10 | AGA (R) | 45 | AGG (R) | 19 |
| CGA (R) | 2 | CGC (R) | 2 | CGG (R) | 3 | CGT (R) | 6 |
| AGC (S) | 12 | AGT (S) | 20 | TCA (S) | 19 | TCC (S) | 2 |
| TCG (S) | 1 | TCT (S) | 22 | ACA (T) | 24 | ACC (T) | 9 |
| ACG (T) | 1 | ACT (T) | 31 | GTA (V) | 21 | GTC (V) | 2 |
| GTG (V) | 2 | GTT (V) | 48 | TGG (W) | 7 | TAC (Y) | 8 |
| TAT (Y) | 47 | TAA (.) | 0 | TGA (.) | 0 | TAG (.) | 0 |

**Comments:** You have inserted a protein sequence from and your optimization criteria have been "Guided random". Your sequence has 1368 AA.

**Abbreviations:** CAI: Codon Adaptation Index. ENc: Effective Number of Codons. %GC: G+C percentage. %AT: A+T percentage

**Supplementary File 2. Concatenated *B. burgdorferi* B31 genome for CRISPy-web.**

B31_genome_contig.gbk

**Supplementary File 3. AlphaFold structure predictions on Google Colab.**

AlphaFold predictions for mature P35G and P35S OspA mutants were superimposed onto the prediction for mature WT *B. burgdorferi* B31 OspA (transparent) and colored by predicted local difference distance test (pLDDT) score for each residue. PyMOL calculated RMSD is shown for each superimposition. Sequences for each prediction are listed in fasta format. Sequences of the mature OspA peptides without signal peptides are shown in Fasta format below. The mutated Pro35 residue is highlighted in blue.

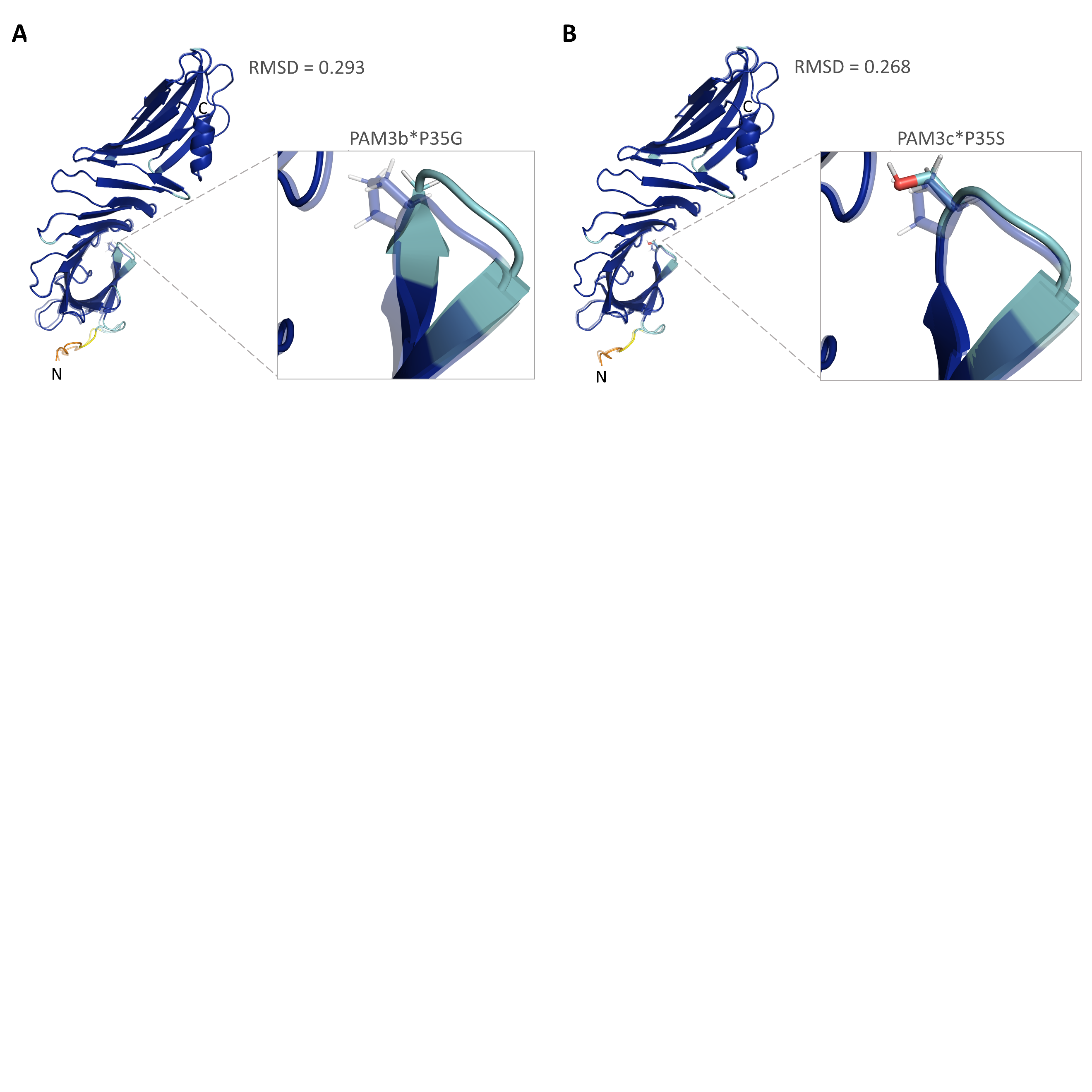

>WT OspA

CKQNVSSLDEKNSVSVDL**P**GEMKVLVSKEKNKDGKYDLIATVDKLELKGTSDKNNGSGVLEGVKADKSKVKLTISDDLGQTTLEVFKEDGKTLVSKKVTSKDKSSTEEKFNEKGEVSEKIITRADGTRLEYTGIKSDGSGKAKEVLKGYVLEGTLTAEKTTLVVKEGTVTLSKNISKSGEVSVELNDTDSSAATKKTAAWNSGTSTLTITVNSKKTKDLVFTKENTITVQQYDSNGTKLEGSAVEITKLDEIKNALK

>OspA Pro35Gly

CKQNVSSLDEKNSVSVDL**G**GEMKVLVSKEKNKDGKYDLIATVDKLELKGTSDKNNGSGVLEGVKADKSKVKLTISDDLGQTTLEVFKEDGKTLVSKKVTSKDKSSTEEKFNEKGEVSEKIITRADGTRLEYTGIKSDGSGKAKEVLKGYVLEGTLTAEKTTLVVKEGTVTLSKNISKSGEVSVELNDTDSSAATKKTAAWNSGTSTLTITVNSKKTKDLVFTKENTITVQQYDSNGTKLEGSAVEITKLDEIKNALK

>OspA Pro35Ser (CA7)

CKQNVSSLDEKNSVSVDL**S**GEMKVLVSKEKNKDGKYDLIATVDKLELKGTSDKNNGSGVLEGVKADKSKVKLTISDDLGQTTLEVFKEDGKTLVSKKVTSKDKSSTEEKFNEKGEVSEKIITRADGTRLEYTGIKSDGSGKAKEVLKGYVLEGTLTAEKTTLVVKEGTVTLSKNISKSGEVSVELNDTDSSAATKKTAAWNSGTSTLTITVNSKKTKDLVFTKENTITVQQYDSNGTKLEGSAVEITKLDEIKNALK
